# Muscle function and homeostasis require macrophage-derived cytokine inhibition of AKT activity in *Drosophila*

**DOI:** 10.1101/763557

**Authors:** Katrin Kierdorf, Fabian Hersperger, Jessica Sharrock, Crystal M. Vincent, Pinar Ustaoglu, Jiawen Dou, Attila Gyoergy, Olaf Groß, Daria E. Siekhaus, Marc S. Dionne

**Author notes:** Current address: Institute of Neuropathology, Faculty of Medicine, University of Freiburg, Breisacherstraße 64, 79106 Freiburg, Germany (Katrin Kierdorf). Current address: Immunology Program, Memorial Sloan-Kettering Cancer Center, New York NY 10065, USA.

## Abstract

*Unpaired* ligands are secreted signals that act via a GP130-like receptor, *domeless*, to activate JAK-STAT signaling in *Drosophila*. Like many mammalian cytokines, *unpaireds* can be activated by infection and other stresses and can promote insulin resistance in target tissues. However, the importance of this effect in non-inflammatory physiology is unknown. Here, we identify a requirement for *unpaired*-JAK signaling as a metabolic regulator in healthy adult *Drosophila* muscle. Adult muscles show basal JAK-STAT signaling activity in the absence of any immune challenge. Macrophages are the source of much of this tonic signal. Loss of the *dome* receptor on adult muscles significantly reduces lifespan and causes local and systemic metabolic pathology. These pathologies result from hyperactivation of AKT and consequent deregulation of metabolism. Thus, we identify a cytokine signal from macrophages to muscle that controls AKT activity and metabolic homeostasis.

## Introduction

JAK/STAT activating signals are critical regulators of many biological processes in animals. Originally described mainly in immune contexts, it has increasingly become clear that JAK/STAT signaling is also central to metabolic regulation in many tissues (Dodington et al., 2018; Villarino et al., 2017). One common consequence of activation of JAK/STAT pathways in inflammatory contexts is insulin resistance in target tissues, including muscle (Kim et al., 2013; Mashili et al., 2013). However, it is difficult to describe a general metabolic interaction between JAK/STAT and insulin signaling in mammals, due to different effects at different developmental stages, differences between acute and chronic actions, and the large number of JAKs and STATs present in mammalian genomes (Dodington et al., 2018; Mavalli et al., 2010; Nieto-Vazquez et al., 2008; Vijayakumar et al., 2013).

The fruit fly *Drosophila melanogaster* has a single, well-conserved JAK-STAT signaling pathway. The *unpaired (upd)* genes *upd1*-*3* encode the three known ligands for this pathway; they signal by binding to a single common GP130-like receptor, encoded by *domeless* (*dome*) (Agaisse et al., 2003; Brown et al., 2001; Chen et al., 2002). Upon ligand binding, the single JAK tyrosine kinase in *Drosophila*, encoded by *hopscotch* (*hop*), is activated; Hop then activates the single known STAT, STAT92E, which functions as a homodimer (Binari and Perrimon, 1994; Chen et al., 2002; Hou et al., 1996; Yan et al., 1996). This signaling pathway plays a wide variety of functions, including segmentation of the early embryo, regulation of hematopoiesis, maintenance and differentiation of stem cells in the gut, and immune modulation (Amoyel and Bach, 2012; Myllymaki and Ramet, 2014). Importantly, several recent studies indicate roles for *upd* cytokines in metabolic regulation; for example, the *upd*s are important nutrient-responsive signals in the adult fly (Beshel et al., 2017; Rajan and Perrimon, 2012; Woodcock et al., 2015; Zhao and Karpac, 2017).

Here, we identify a physiological requirement for Dome signaling in adult muscle. We observe that adult muscles show significant JAK/STAT signaling activity in the absence of obvious immune challenge and macrophages seem to be a source of this signal. Inactivation of *dome* on adult muscles significantly reduces lifespan and causes muscular pathology and physiological dysfunction; these result from remarkably strong AKT hyperactivation and consequent dysregulation of metabolism. We thus describe a new role for JAK/STAT signaling in adult *Drosophila* muscle with critical importance in healthy metabolic regulation.

## Results

### *dome* is required in adult muscle

To find physiological functions of JAK/STAT signaling in the adult fly, we identified tissues with basal JAK/STAT pathway activity using a STAT-responsive GFP reporter (*10xSTAT92E-GFP*) (Bach et al., 2007). The strongest reporter activity we observed was in legs and thorax. We examined flies also carrying a muscle myosin heavy chain RFP reporter (*MHC-RFP*) and observed co-localization of GFP and RFP expression in the muscles of the legs, thorax and body wall (Fig S1A). We observed strong, somewhat heterogeneous reporter expression in all the muscles of the thorax and the legs, with strong expression in various leg and jump muscles and apparently weaker expression throughout the body wall muscles and indirect flight muscles (Fig 1A).

**Figure 1.**
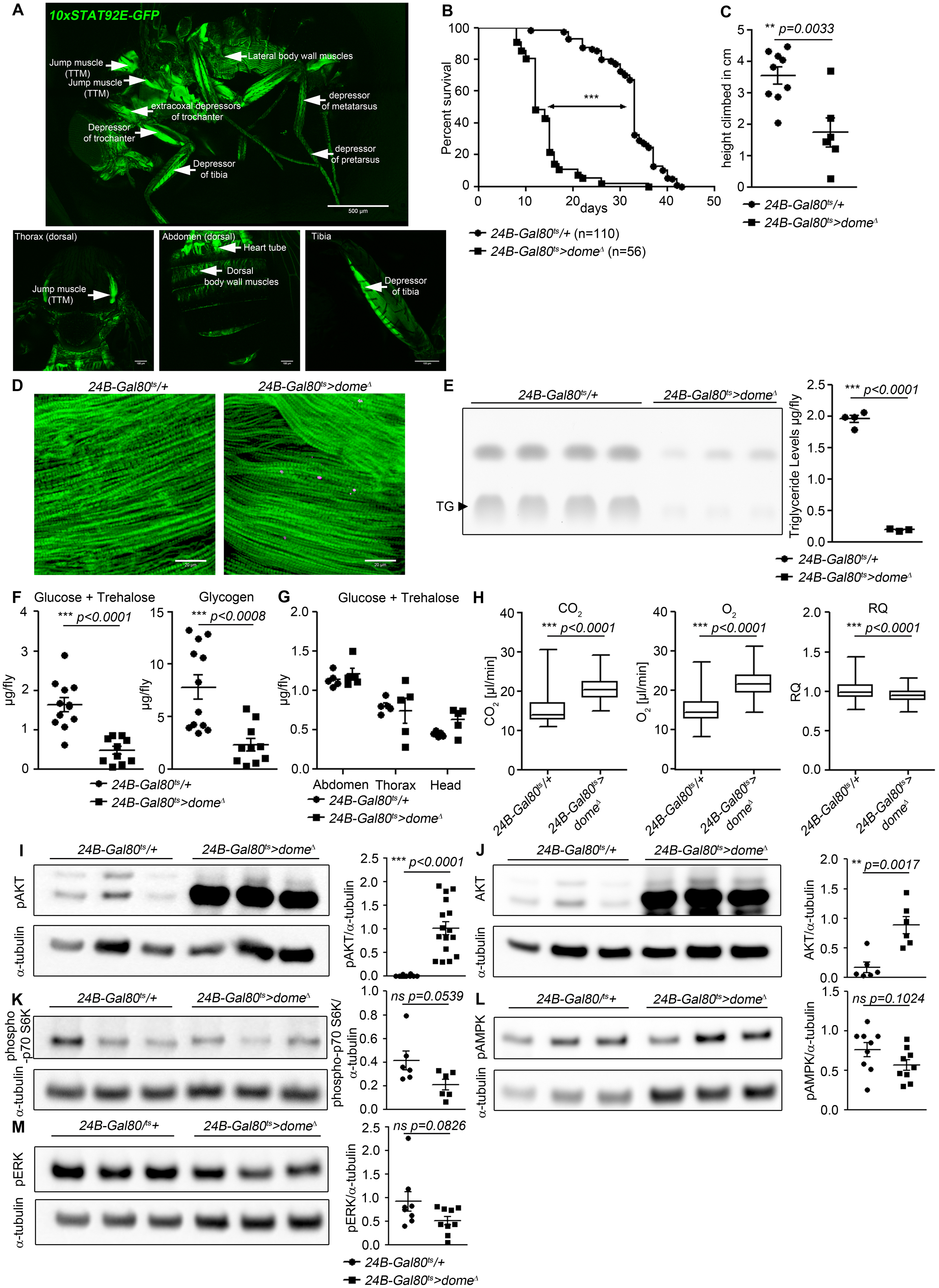
Dome inhibition in adult muscle reduces lifespan, disrupts homeostasis, and causes AKT hyperactivation. (A) STAT activity in different muscles in 10xSTAT92E-GFP reporter fly. One fly out of 5 shown. Upper panel: lateral view, Scale bar=500μm. Lower panels: dorsal thorax (left); dorsal abdomen (middle); tibia (right), Scale bar=100μm (B) Lifespan of *24B-Gal80*^*ts*^*/+* and *24B-Gal80*^*ts*^>*dome*^*Δ*^ at 29°, pooled from three independent experiments. Log-Rank test: χ^2^ =166, *** p<0.0001; Wilcoxon test: χ^2^ =157.7, *** p<0.0001. (C) Negative geotaxis assay of 14-day-old *24B-Gal80*^*ts*^*/+* and *24B-Gal80*^*ts*^>*dome*^*Δ*^ flies. Points represent mean height climbed in individual vials (~20 flies/vial), pooled from three independent experiments. (D) Muscle (Phalloidin) and neutral lipid (LipidTox) of thorax samples from 14-day-old *24B-Gal80*^*ts*^*/+* and *24B-Gal80*^*ts*^>*dome*^*Δ*^ flies. One representative fly per genotype is shown of six analysed. Scale bar=50μm. (E) Thin layer chromatography (TLC) of triglycerides in 7-day-old *24B-Gal80*^*ts*^*/+* and *24B-Gal80*^*ts*^>*dome*^*Δ*^ flies, n=3-4 per genotype. One experiment of two is shown. (F) Glucose and trehalose (left) and glycogen (right) in 7-day-old *24B-Gal80*^*ts*^*/+* and *24B-Gal80*^*ts*^>*dome*^*Δ*^ flies, pooled from two independent experiments. (G) Glucose and trehalose content of dissected abdomen, thorax, and head of 7-day-old *24B-Gal80*^*ts*^*/+* and *24B-Gal80*^*ts*^>*dome*^*Δ*^ flies. (H) CO_2_ produced, O_2_ consumed, and RQ of 7-day-old *24B-Gal80*^*ts*^*/+* and *24B-Gal80^ts^>dome^Δ^* flies. Box plots show data from one representative experiment of three, with data collected from a 24 h measurement pooled from 3-4 tubes per genotype with 10 flies/tube. P values from Mann-Whitney test. (I-M) Western blots of leg protein from 14-day-old *24B-Gal80*^*ts*^*/+* and *24B-Gal80*^*ts*^>*dome*^*Δ*^ flies. (I) Phospho-AKT (S505). One experiment of four is shown. (J) Total AKT. One experiment of two is shown. (K) Phospho-p70 S6K (T398). One experiment of two is shown. (L) Phospho-AMPKα (T173). One experiment of three is shown. (M) Phospho-ERK (T202/Y204). One experiment of three is shown. P values in C, E, F, I-M from unpaired T-test.

*dome* encodes the only known *Drosophila* STAT-activating receptor. To investigate the physiological role of this signal, we expressed *dome*^*Δ*^, a dominant-negative version of Dome lacking the intracellular signaling domain, with a temperature-inducible muscle specific driver line (*w;tubulin-Gal80*^*ts*^;*24B-Gal4*) (Fig S1B) (Brown et al., 2001). Controls (*24B-Gal80*^*ts*^*/+*) and experimental flies (*24B-Gal80*^*ts*^>*dome*^*Δ*^) were raised at 18° until eclosion to permit Dome activity during development. Flies were then shifted to 29° to inhibit Dome activity and their lifespan was monitored. Flies with Dome signaling inhibited in adult muscles were short-lived (Fig 1B, Fig S1C). This effect was also observed, more weakly, in flies kept at 25° (Fig S1D). Upd-JAK-STAT signaling is important to maintain gut integrity, and defects in gut integrity often precede death in *Drosophil*a; however, our flies did not exhibit loss of gut integrity (Fig S1E) (Jiang et al., 2009; Rera et al., 2012). To determine whether Dome inhibition caused meaningful physiological dysfunction, we assayed climbing activity in *24B-Gal80*^*ts*^*/+* control flies and *24B-Gal80*^*ts*^>*dome*^*Δ*^ flies. *24B-Gal80*^*ts*^>*dome*^*Δ*^ flies showed significantly impaired climbing compared to controls (Fig 1C). Adult muscle-specific expression of *dome*^*Δ*^ with a second Gal4 line (*w;tub-Gal80*^*ts*^;*Mef2-Gal4*) gave a similar reduction in lifespan and decline in climbing activity, confirming that the defect resulted from a requirement for Dome activity in muscle (Fig S1F, G).

Impaired muscle function is sometimes accompanied by lipid accumulation (Baik et al., 2017). Therefore, we stained thorax muscles with the neutral lipid dye LipidTox. In 14 day old flies, we detected numerous small neutral lipid inclusions in several muscles, including the large jump muscle (TTM), of *24B-Gal80*^*ts*^>*dome*^*Δ*^ flies (Fig 1D).

### Muscle *dome* activity is required for normal systemic homeostasis

Having observed lipid inclusions in adult muscles, we analysed the systemic metabolic state of *24B-Gal80*^*ts*^>*dome*^*Δ*^ flies. We observed significant reductions in total triglyceride, glycogen and free sugar (glucose + trehalose) in these animals (Fig 1E, F). The reduction in free sugar was not detectable in any dissected solid tissue, suggesting that it was due to a reduction in hemolymph sugar (Fig 1G).

Reduced hemolymph sugar could result from increased tissue glucose uptake. In this case, it should be reflected in an increased metabolic stores or metabolic rate. Since metabolic stores were decreased in our flies, we tested metabolic rate by measuring respiration. CO_2_ production and O_2_ consumption were both significantly increased in *24B-Gal80*^*ts*^>*dome*^*Δ*^ flies, indicating an overall increase in metabolic rate (Fig 1H).

### *dome* acts via *hop* to regulate AKT activity with little effect on other nutrient signaling pathways

The observed metabolic changes imply differences in activity of nutrient-regulated signaling pathways in *24B-Gal80*^*ts*^>*dome*^*Δ*^ flies. Several signaling pathways respond to nutrients, or their absence, to coordinate energy consumption and storage (Britton et al., 2002; Lizcano et al., 2003; Ulgherait et al., 2014). Of these, insulin signaling via AKT is the primary driver of sugar uptake by peripheral tissues.

We examined the activity of these signaling mechanisms in legs (a tissue source strongly enriched in muscle) from *24B-Gal80*^*ts*^>*dome*^*Δ*^ flies. We found an extremely strong increase in abundance of the 60-kDa form of total and activated (S505-phosphorylated) AKT (Fig 1I, J). This change was also seen in legs from *Mef2-Gal80*^*ts*^>*dome*^*Δ*^ flies, confirming that *dome* functions in muscles (Fig S1H, I). We also saw this effect in flies carrying a different insertion of the *dome*^*Δ*^ transgene, under the control of a third muscle-specific driver, *MHC-Gal4*, though the effect was weaker (Fig S1J). These *MHC-Gal4*>*dome*^*Δ*^*(II)* animals were also short-lived relative to controls (Fig S1K).

Elevated total AKT could result from increased transcript abundance or changes in protein production or stability. We distinguished between these possibilities by assaying *Akt1* mRNA; *Akt1* transcript levels were elevated in *24B-Gal80*^*ts*^>*dome*^*Δ*^ muscle, but only by about 75%, suggesting that the large effect on AKT protein abundance must be, at least in part, post-transcriptional (Fig S1L). Similarly, AKT hyperactivation could be driven by insulin-like peptide overexpression; however, we assayed the expression of *Ilp2-7* in whole flies and observed that none of these peptides were significantly overexpressed (Fig S1M-R).

Unlike AKT, the amino-acid-responsive TORC1/S6K and the starvation-responsive AMPK pathway showed no significant difference in activity in *24B-Gal80*^*ts*^>*dome*^*Δ*^ flies (Fig 1K, L). However, flies with AMPK knocked down in muscle did exhibit mild AKT hyperactivation (Fig S2A).

To identify signaling mediators acting between Dome and AKT, we first tested activity of the MAPK-ERK pathway, which can act downstream of the JAK kinase Hop (Luo et al., 2002). We found an insignificant reduction in ERK activity in *24B-Gal80*^*ts*^>*dome*^*Δ*^ flies (Fig 1M). We then assayed survival and AKT activity in flies with *hop* (JAK), *Dsor1* (MEK) and *rl* (ERK) knocked down in adult muscle. *rl* and *Dsor1* knockdown gave mild or no effect on survival and pAKT (Fig S2B, C). In contrast, *hop* knockdown phenocopied the milder *dome*^*Δ*^ transgene with regard to survival and pAKT (Fig S2D, E).

We further analysed the requirement for *hop* in muscle *dome* signaling by placing *24B-Gal80*^*ts*^>*dome*^*Δ*^ on a genetic background carrying the viable gain-of-function allele *hop*^*Tum-l*^. Flies carrying *hop*^*Tum-l*^ alone exhibited no change in lifespan, AKT phosphorylation, or muscle lipid deposition (Fig 2A-C). However, *hop*^*Tum-l*^ completely rescued lifespan and pAKT levels in *24B-Gal80*^*ts*^>*dome*^*Δ*^ flies (Fig 2D, E), indicating that the physiological activity of muscle Dome is mediated via Hop and that this signal is required, but not sufficient, to control muscle AKT activity.

**Figure 2.**
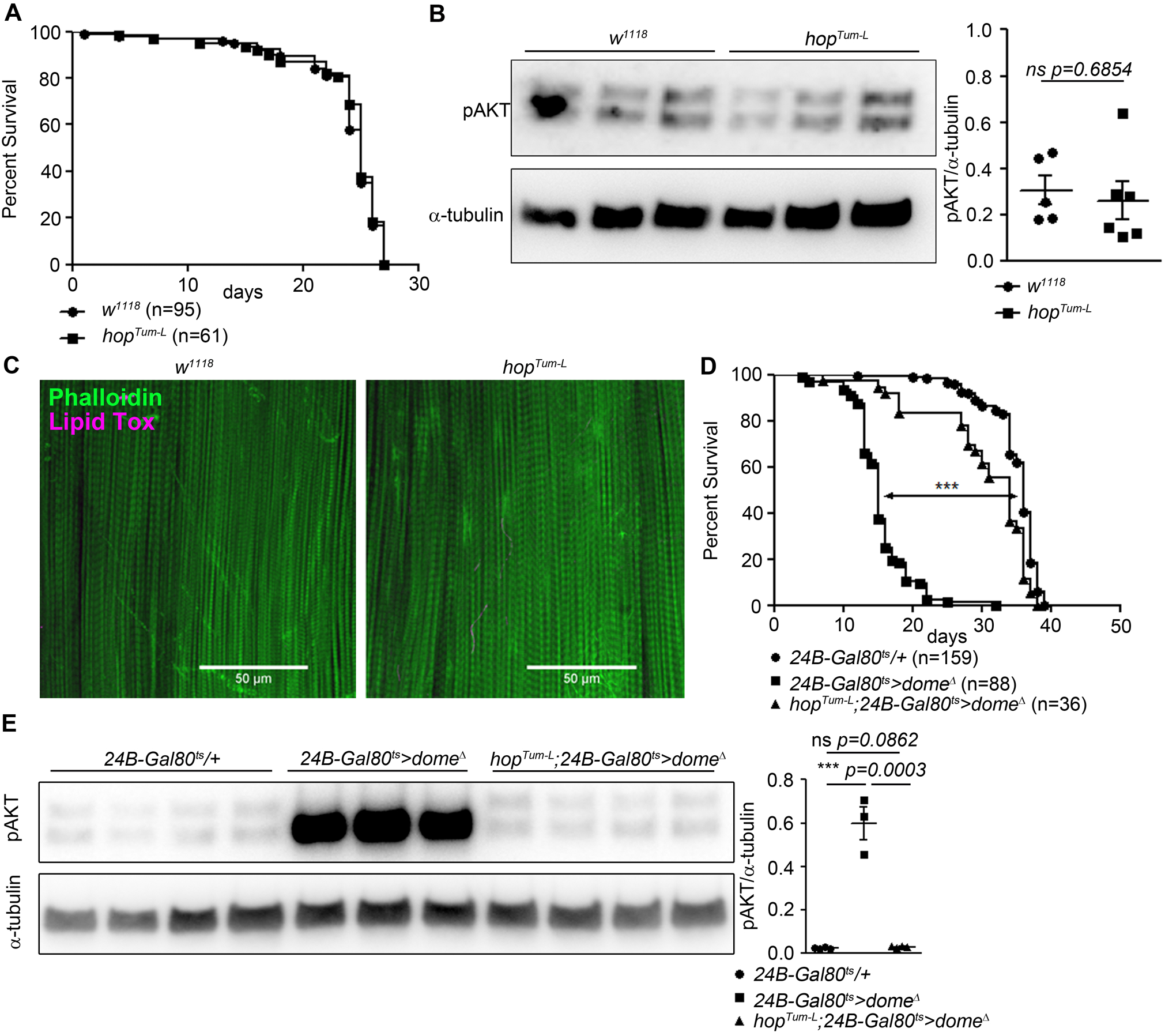
Hop is required, but not sufficient, for Dome to control AKT. (A) Lifespan of *w*^*1118*^ and *hop*^*Tum-l*^ flies at 29°, pooled from two independent experiments. Log-Rank test: χ^2^ =0.3223, ns p=0.5702; Wilcoxon test: χ^2^ =0.4756, ns p=0.4906. (B) Phospho-AKT in leg samples from 14-day-old *w*^*1118*^ and *hop*^*Tum-l*^ flies. One experiment of two is shown. (C) Actin (Phalloidin) and neutral lipid (LipidTox) in flight muscle from 14-day-old *w*^*1118*^ and *hop*^*Tum-l*^ flies. One representative fly shown of six analysed per genotype. Scale bar=50μm. (D) Lifespan of *24B-Gal80*^*ts*^*/+*, *24B-Gal80*^*ts*^>*dome*^*Δ*^, and *hop*^*Tum-L*^;*24B-Gal80*^*ts*^>*dome*^*Δ*^ flies at 29°, pooled from four independent experiments. Log-Rank test (*24B-Gal80*^*ts*^*/+* vs. *24B-Gal80*^*ts*^>*dome*^*Δ*^): χ^2^ =319.4, *** p<0.0001; Wilcoxon test (*24B-Gal80*^*ts*^*/+* vs. *24B-Gal80*^*ts*^>*dome*^*Δ*^): χ^2^ =280.2, *** p<0.0001. Log-Rank test (*24B-Gal80*^*ts*^*/+* vs. *hop*^*Tum-L*^ *24B-Gal80*^*ts*^>*dome*^*Δ*^): χ^2^ =18.87, *** p<0.0001; Wilcoxon test (*24B-Gal80*^*ts*^*/+* vs. *hop*^*Tum-L*^ *24B-Gal80*^*ts*^>*dome*^*Δ*^): χ^2^ =20.83, *** p<0.0001. (E) Phospho-AKT in leg samples from 14-day-old *24B-Gal80*^*ts*^*/+*, *24B-Gal80*^*ts*^>*dome*^*Δ*^ and *hop*^*Tum-L*^;*24B-Gal80*^*ts*^>*dome*^*Δ*^ flies. P values in B, E from unpaired T-test.

### Increased AKT activity causes the effects of *dome* inhibition

The phenotype of *24B-Gal80*^*ts*^>*dome*^*Δ*^ flies is similar to that previously described in flies with loss of function in *Pten* or *foxo* (Demontis and Perrimon, 2010; Mensah et al., 2015), suggesting that AKT hyperactivation might cause the *dome* loss of function phenotype; however, to our knowledge, direct activation of muscle AKT had not previously been analysed. We generated flies with inducible expression of activated AKT (*myr-AKT*) in adult muscles (*w;tubulin-Gal80*^*ts*^*/+;24B-Gal4/UAS-myr-AKT* [*24B-Gal80*^*ts*^>*myr-AKT*]) (Stocker et al., 2002). These animals phenocopied *24B-Gal80*^*ts*^>*dome*^*Δ*^ flies with regard to lifespan, climbing activity, metabolite levels, metabolic rate, and muscle lipid deposition (Fig 3A-F).

**Figure 3.**
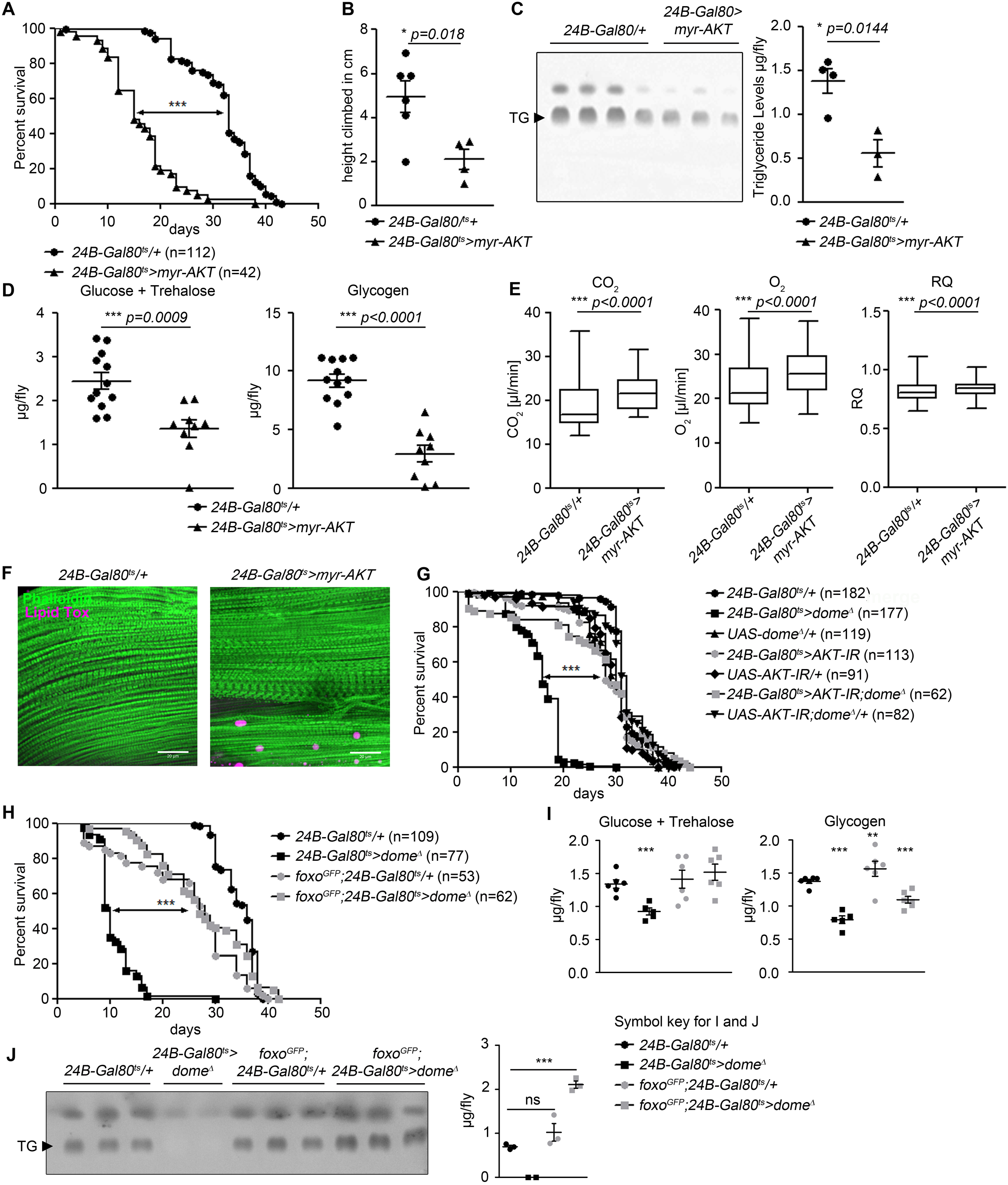
AKT hyperactivation causes pathology in *24B-Gal80*^*ts*^>*dome*^*Δ*^ flies. (A) Lifespan of *24B-Gal80*^*ts*^*/+* and *24B-Gal80^ts^>myr-AKT* at 29°, pooled data from three independent experiments. Log-Rank test: χ^2^ =115.5, *** p<0.0001; Wilcoxon test: χ^2^ =123.6, *** p<0.0001. (B) Negative geotaxis assay of 14-day-old *24B-Gal80*^*ts*^*/+* and *24B-Gal80^ts^>myr-AKT* flies. Points represent mean height climbed in individual vials (~20 flies/vial), pooled from two independent experiments. (C) TLC of triglycerides in 7-day-old *24B-Gal80*^*ts*^*/+* and *24B-Gal80*^*ts*^>*myr-AKT* flies, n=3-4 per genotype. One experiment of two is shown. (D) Glucose and trehalose (left panel) and glycogen (right panel) in 7-day-old *24B-Gal80*^*ts*^*/+* (n=12) and *24B-Gal80*^*ts*^>*myr-AKT* (n=9) flies, pooled from two independent experiments. (E) CO_2_ produced, O_2_ consumed, and RQ of 7-day-old *24B-Gal80*^*ts*^*/+* and *24B-Gal80*^*ts*^>*myr-AKT* flies. Box plots show data from one representative experiment of three, with data points collected from a 24 h measurement pooled from 3-4 tubes per genotype with 10 flies/tube. P values from Mann-Whitney test. (F) Phalloidin and LipidTox staining of thorax samples from 14-day-old *24B-Gal80*^*ts*^*/+* and *24B-Gal80*^*ts*^>*myr-AKT* flies. One representative fly per genotype is shown of 3 analysed per group in 2 independent experiments. Scale bar=50μm. (G) Lifespan of *24B-Gal80*^*ts*^*/+*, *24B-Gal80*^*ts*^>*dome*^*Δ*^, *UAS-dome^Δ^/+, 24B-Gal80^ts^>AKT-IR*, *UAS-AKT-IR/+*, *24B-Gal80*^*ts*^>*AKT-IR;dome*^*Δ*^ and UAS-*AKT-IR;dome^Δ^/+* flies at 29°. One from four independent experiments shown. Log-Rank test (*24B-Gal80*^*ts*^>*dome*^*Δ*^ vs. *24B-Gal80^ts^>AKT-IR;dome^Δ^*): χ^2^ =101.0, *** p<0.0001; Wilcoxon test (*24B-Gal80*^*ts*^>*dome*^*Δ*^ vs. *24B-Gal80^ts^>AKT-IR;dome^Δ^*): χ^2^ =59.87, *** p<0.0001. (H) Lifespan of *24B-Gal80*^*ts*^*/+*, *24B-Gal80*^*ts*^>*dome*^*Δ*^, *foxo-GFP;24B-Gal80^ts^/+*, and *foxo-GFP;24B-Gal80*^*ts*^>*dome*^*Δ*^ flies at 29°, pooled from three independent experiments. Log-Rank test (*24B-Gal80*^*ts*^>*dome*^*Δ*^ vs. *foxo-GFP;24B-Gal80*^*ts*^>*dome*^*Δ*^): χ^2^ =114.0, *** p<0.0001; Wilcoxon test (*24B-Gal80*^*ts*^>*dome*^*Δ*^ vs. *foxo-GFP;24B-Gal80^ts^>dome^Δ^*): χ^2^ =93.59, *** p<0.0001. *24B-Gal80*^*ts*^*/+* and *24B-Gal80*^*ts*^>*dome*^*Δ*^ controls in G and H are the same because a single survival experiment was split into two graphs. (I) Glucose + trehalose and glycogen in 7-day-old *24B-Gal80*^*ts*^*/+*, *24B-Gal80*^*ts*^>*dome*^*Δ*^, *foxo-GFP;24B-Gal80/+*, and foxo-GFP; *24B-Gal80*^*ts*^>*dome*^*Δ*^ flies. (J) TLC of triglycerides in 7-day-old *24B-Gal80*^*ts*^*/+*, *24B-Gal80*^*ts*^>*dome*^*Δ*^, *foxo-GFP;24B-Gal80^ts^/+*, and *foxo-GFP;24B-Gal80*^*ts*^*dome*^*Δ*^ flies. P values in B-D, I, J from unpaired T-test.

We concluded that AKT hyperactivation could cause the pathologies seen in *24B-Gal80*^*ts*^>*dome*^*Δ*^ flies. We next tested whether reducing AKT activity could rescue *24B-Gal80*^*ts*^>*dome*^*Δ*^ flies. We generated flies carrying muscle-specific inducible dominant negative dome (*UAS-dome*^*Δ*^) with dsRNA against *Akt1* (*UAS-AKT-IR*). These flies showed significantly longer lifespan than *24B-Gal80*^*ts*^>*dome*^*Δ*^ and *24B-Gal80*^*ts*^>*AKT-IR* flies, similar to all control genotypes analyzed (Fig 3G). Dome and AKT antagonism synergised to control the mRNA level of *dome* itself, further suggesting strong mutual antagonism between these pathways (Fig S3A).

AKT hyperactivation should reduce FOXO transcriptional activity. To test whether this loss of FOXO activity caused some of the pathologies observed in *24B-Gal80*^*ts*^>*dome*^*Δ*^ flies, we increased *foxo* gene dosage by combining *24B-Gal80*^*ts*^>*dome*^*Δ*^ with a transgene carrying a FOXO-GFP fusion protein under the control of the endogenous *foxo* regulatory regions. These animals exhibited rescue of physiological defects and lifespan compared to *24B-Gal80*^*ts*^>*dome*^*Δ*^ flies (Fig 3H-J). They also exhibited increased *dome* expression (Fig S3B). The effects of these manipulations on published *foxo* target genes were mixed (Fig S3B); the strongest effect we observed was that Dome blockade increased *upd2* expression, consistent with the observation that FOXO activity inhibits *upd2* expression in muscle (none of the other genes tested have been shown to be FOXO targets in muscle) (Zhao and Karpac, 2017). This may explain some of the systemic effects of Dome blockade.

The effect of the *foxo* transgene was stronger than expected from a 1.5-fold increase in *foxo* expression, so we further explored the relationship between FOXO protein expression and AKT phosphorylation. We found that *24B-Gal80*^*ts*^>*dome*^*Δ*^ markedly increased FOXO-GFP abundance, so that the increase in total FOXO was much greater than 1.5-fold (Fig S3C). This drove an apparent feedback effect, restoring AKT in leg samples of *foxo*^*GFP*^;*24B-Gal80*^*ts*^>*dome*^*Δ*^ flies to near-normal levels (Fig S3D).

### Macrophages are a relevant source of *upd* signals

Plasmatocytes—*Drosophila* macrophages—are a key source of *upd3* in flies on high fat diet and in mycobacterial infection (Péan et al., 2017; Woodcock et al., 2015). Plasmatocytes also express *upd1-3* in unchallenged flies (Chakrabarti et al., 2016). We thus tested their role in activation of muscle Dome.

We found plasmatocytes close to STAT-GFP-positive leg muscle (Fig 4A, B). This, and the prior published data, suggested that plasmatocytes might produce relevant levels of *dome-*activating cytokines in steady state. We then overexpressed *upd3* in plasmatocytes and observed a potent increase in muscle STAT-GFP activity (Fig 4C), confirming that plasmatocyte-derived *upd* signals were able to activate muscle Dome.

**Figure 4.**
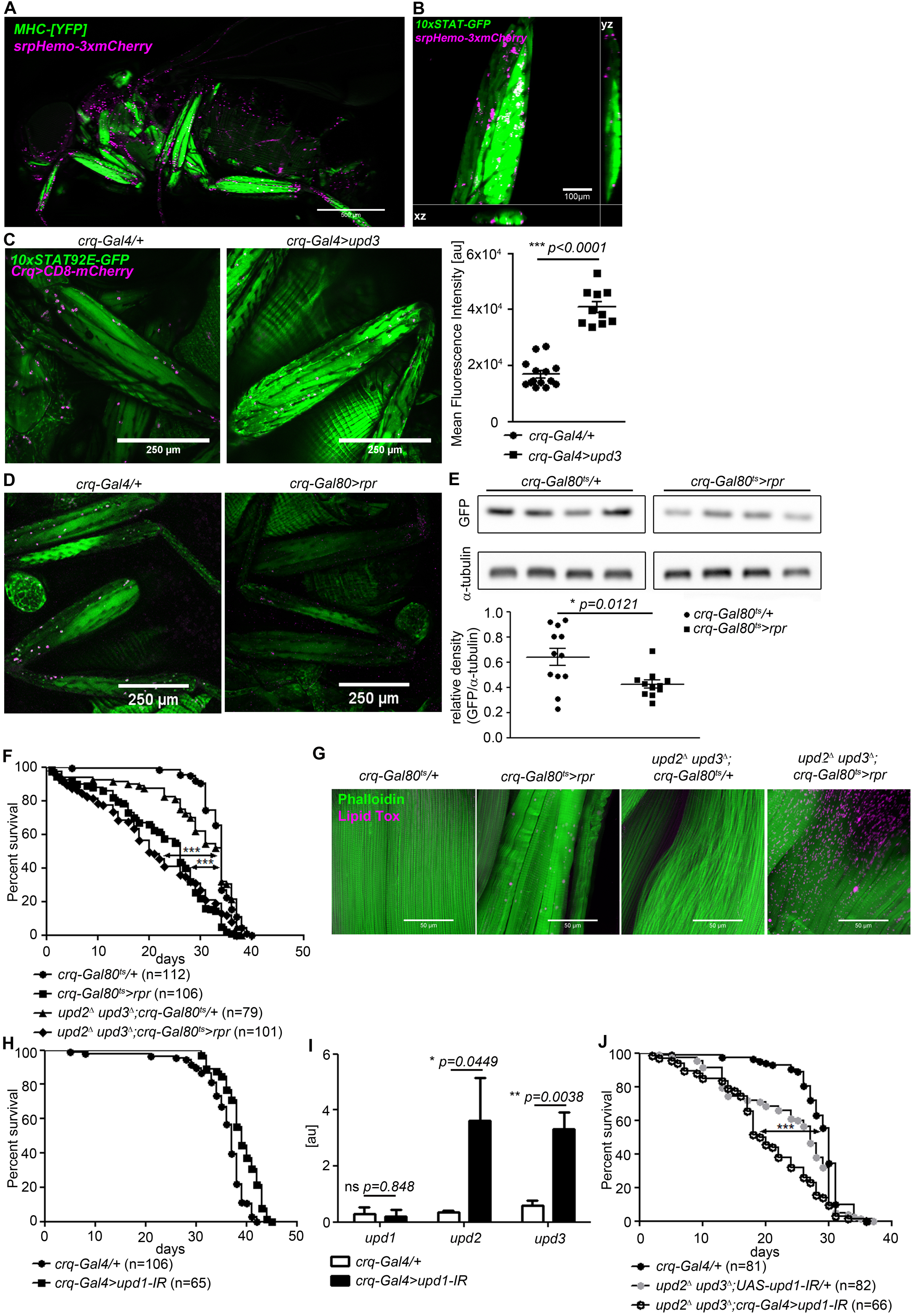
Plasmatocytes promote muscle Dome activity. (A) Muscle (*MHC^YFP^*) and plasmatocytes (*srpHemo-3xmCherry*) in 7-day-old flies. Plasmatocytes are found in close proximity to adult muscles. One representative fly of 5 is shown. Scale bar=500μm. (B) Legs and plasmatocytes in 7-day-old *10xSTAT92E-GFP;srpHemo-3xmCherry* flies. Muscle with high JAK-STAT activity (green) is surrounded by plasmatocytes (magenta). One representative fly of 5 is shown. Scale bar=100μm. (C) STAT activity and plasmatocytes in legs from control (*10xSTAT92E-GFP;crq-Gal4>CD8-mCherry/+*) and *upd3*-overexpressing (*10xSTAT92E-GFP;crq-4>CD8mCherry/UAS-upd3*) flies. One representative fly of 10-14 is shown. Scale bar=100μm. Graph shows mean fluorescence intensity (MFI). (D) STAT activity and plasmatocytes in legs from control (*10xSTAT92E-GFP;crq-Gal80^ts^>CD8-mCherry/+*) and plasmatocyte-depleted (*10xSTAT92E-GFP;crq-Gal80^ts^>CD8mCherry/rpr*) flies. One representative fly of six is shown. Scale bar=250μm. (E) Western blot analysis of STAT-driven GFP in legs from 7-day-old control (*10xSTAT92E-GFP;crq-Gal80^ts^>CD8-mCherry/+*) and plasmatocyte-depleted (*10xSTAT92E-GFP;crq-Gal80^ts^>CD8-mCherry/rpr* flies). One representative experiment of three is shown. Graph shows STAT-GFP/α-tubulin for control (*crq-Gal80*^*ts*^*/+*) and plasmatocyte-depleted (*crq-Gal80^ts^>rpr*) leg samples. (F) Lifespan of *crq-Gal80*^*ts*^*/+*, *crq-Gal80^ts^>rpr*, *upd2^Δ^ upd3^Δ^;crq-Gal80^ts^/+*, and *upd2^Δ^ upd3^Δ^;crq-Gal80^ts^>rpr* flies at 29°; pooled data from three independent experiments shown. Log-Rank test (*crq-Gal80*^*ts*^*/+* vs. *crq-Gal80^ts^>rpr*): χ2 =101.7, *** p<0.0001; Wilcoxon test (*crq-Gal80*^*ts*^*/+* vs. *crq-Gal80^ts^>rpr*): χ2 =107.8, *** p<0.0001; Log-Rank test (*crq-Gal80*^*ts*^*/+* vs. *upd2 ^Δ^ upd3^Δ^;crq-Gal80^ts^>rpr*): χ2 =60.03, *** p<0.0001; Wilcoxon test (*crq-Gal80*^*ts*^*/+* vs. *upd2 ^Δ^ upd3^Δ^;crq-Gal80^ts^>rpr*): χ2 =80.97, *** p<0.0001. (G) Actin (Phalloidin) and neutral lipid (LipidTox) in thorax samples from 14-day-old *crq-Gal80*^*ts*^*/+*, *crq-Gal80^ts^>rpr*, *upd2 ^Δ^ upd3^Δ^;crq-Gal80^ts^/+*, and *upd2 ^Δ^ upd3^Δ^;crq-Gal80^ts^>rpr* flies. One representative fly per genotype shown of 6 analysed per group. Scale bar=50μm. (H) Lifespan of *crq-Gal4/+* and *crq-Gal4>upd1-IR* flies at 29°. Log-Rank test: χ2 =31.36, *** p<0.0001; Wilcoxon test: χ2 =22.17, *** p=0.0001. (I) Expression by qRT-PCR of *upd1*, *upd2* and *upd3* in thorax samples of *crq-Gal4/+* and *crq-Gal4>upd1-IR* flies, data from four independent samples of each genotype. (J) Lifespan of *crq-Gal4/+*, *upd2 ^Δ^ upd3^Δ^;UAS-upd1-IR/+*, and *upd2^Δ^ upd3^Δ^;crq-Gal4>upd1-IR* flies at 29°. Pooled data from three independent experiments shown. Log-Rank test (*crq-Gal4/+* vs. *upd2 ^Δ^ upd3^Δ^;crq-Gal4>upd1-IR*): χ2 =41.12, *** p<0.0001; Wilcoxon test (*crq-Gal4/+* vs. *upd2^Δ^ upd3^Δ^;crq-Gal4>upd1-IR*): χ2 =54.47, *** p<0.0001 Log-Rank test (*crq-Gal4/+* vs. *upd2 ^Δ^ upd3^Δ^;UAS-upd1-IR/+*): χ2 =14.46, *** p<0.0001; Wilcoxon test (*crq-Gal4/+* vs. *upd2^Δ^ upd3^Δ^;UAS-upd1-IR/+*): χ2 =19.99, *** p<0.0001. P values in C, E, H from unpaired T-test.

To determine the physiological relevance of plasmatocyte-derived signals, we assayed STAT-GFP activity in flies in which plasmatocytes had been depleted by expression of the pro-apoptotic gene *reape*r (*rpr*) using a temperature-inducible plasmatocyte-specific driver line (*w;tub-Gal80*^*ts*^;*crq-Gal4*). STAT-GFP fluorescence and GFP abundance were reduced in legs of plasmatocyte-depleted flies (*crq-Gal80*^*ts*^>*rpr*) compared to controls (*crq-Gal80*^*ts*^*/+*) (Fig 4D, E). Activity was not eliminated, indicating that plasmatocytes are not the only source of muscle STAT-activating signals.

We then examined the lifespan of flies in which we had depleted plasmatocytes in combination with various *upd* mutations and knockdowns. Plasmatocyte depletion gave animals that were short-lived (Fig 4F). (This effect was different from that we previously reported, possibly due to changes in fly culture associated with an intervening laboratory move (Woodcock et al., 2015).) The lifespan of these animals was further reduced by combining plasmatocyte depletion with null mutations in *upd2* and *upd3*; plasmatocyte-replete *upd2 upd3* mutants exhibited near-normal lifespan (Fig 4F). Similarly, plasmatocyte depletion drove muscle lipid accumulation, and *upd2 upd3* mutation synergised with plasmatocyte depletion to further increase muscle lipid (Fig 4G). However, depleting plasmatocytes in *upd2 upd3* mutants failed to recapitulate the effects of muscle Dome inhibition on whole-animal triglyceride, free sugar, and glycogen levels (Fig S4A, B). This could be due to antagonistic effects of other plasmatocyte-derived signals.

We attempted to pinpoint a specific Upd as the relevant physiological ligand by examining STAT-GFP activity, first testing mutants in *upd2* and *upd3* because *upd1* mutation is lethal. However, these mutants, including the *upd2 upd3* double-mutant, were apparently normal (Fig S4C). We then tested plasmatocyte-specific knockdown of *upd1* and *upd3*; these animals were also essentially normal (Fig S4D), and plasmatocyte *upd1* knockdown did not reduce lifespan (Fig 4H). However, plasmatocyte-specific *upd1* knockdown gave significant compensating increases in expression of *upd2* and *upd3* (Fig 4I). In keeping with this, combining plasmatocyte-specific *upd1* knockdown with mutations in *upd2* and *upd3* reduced lifespan (Fig 4J) and also reduced STAT-GFP activity in these flies (Fig S4F).

Our results indicate that plasmatocytes are an important physiological source of the Upd signal driving muscle Dome activity in healthy flies, and suggest that *upd1* may be the primary relevant signal in healthy animals. However, plasmatocytes are not the only relevant source of signal, and Upd mutual regulation prevents us from pinpointing a single responsible signal.

## Discussion

Here we show that *upd-dome* signaling in muscle acts via AKT to regulate physiological homeostasis in *Drosophila*. Loss of Dome activity in adult muscles shortens lifespan and promotes local and systemic metabolic disruption. Dome specifically regulates the level and activity of AKT; AKT hyper-activation mediates the observed pathology. Plasmatocytes are a primary source of the cytokine signal. In healthy adult flies, insulin-like peptides are the primary physiological AKT agonists. The effect we observe thus appears to be an example of a cytokine-Dome-JAK signal that impairs insulin function to permit healthy physiology.

Our work fits into a recent body of literature demonstrating key physiological roles for JAK-STAT activating signals in *Drosophila*. Upd1 acts locally in the brain to regulate feeding and energy storage by altering the secretion of neuropeptide F (NPF) (Beshel et al., 2017). Upd2 is released by the fat body in response to dietary triglyceride and sugar to regulate secretion of insulin-like peptides (Rajan and Perrimon, 2012). More recently, muscle-derived Upd2, under control of FOXO, has been shown to regulate production of the glucagon-like signal Akh (Zhao and Karpac, 2017). Indeed, we observe that *upd2* is upregulated in flies with Dome signaling blocked in muscle, possibly explaining some of the systemic metabolic effects we observe. Plasmatocyte-derived Upd3 in flies on a high fat diet can activate the JAK/STAT pathway in various organs including muscles and can promote insulin insensitivity (Woodcock et al., 2015). Our observation that Upd signaling is required to control AKT accumulation and thus insulin pathway activity in healthy adult muscle may explain some of these prior observations and reveals a new role for macrophage-derived cytokine signaling in healthy metabolic regulation.

Several recent reports have examined roles of JAK/STAT signaling in *Drosophila* muscle. In larvae, muscle JAK/STAT signaling can have an effect opposite to the one we report, with pathway loss of function resulting in reduced AKT activity (Yang and Hultmark, 2017). It is unclear whether this difference represents a difference in function between developmental stages (larva vs adult) or a difference between acute and chronic consequences of pathway inactivation. Roles in specific muscle populations have also been described: for example, JAK/STAT signaling in adult visceral muscle regulates expression of Vein, an EGF-family ligand, to control intestinal stem cell proliferation (Buchon et al., 2010; Jiang et al., 2011); the role of this system in other muscles may be analogous, controlling expression of various signals to regulate systemic physiology.

The roles of mammalian JAK/STAT signaling in muscle physiology are more complex, but exhibit several parallels with the fly. In mice, early muscle-specific deletion of Growth Hormone Receptor (GHR) causes several symptoms including insulin resistance, while adult muscle-specific GHR deletion causes entirely different effects, including increased metabolic rate and insulin sensitivity on a high-fat diet (Mavalli et al., 2010; Vijayakumar et al., 2013; Vijayakumar et al., 2012). GHR signals via STAT5; STAT5 deletion in adult skeletal muscle promotes muscle lipid accumulation on a high-fat diet (Baik et al., 2017). Other STAT pathways can also play roles. For example, the JAK-STAT activating cytokine IL-6, which signals primarily via STAT3, increases skeletal muscle insulin sensitivity when given acutely but can drive insulin resistance when provided chronically (Nieto-Vazquez et al., 2008). STAT3 itself can promote muscle insulin resistance (Kim et al., 2013; Mashili et al., 2013). The relationship between these effects and those we have shown here, and the mechanisms regulating plasmatocyte Upd production during healthy physiology, remain to be determined.

## Materials and methods

### Drosophila melanogaster stocks and culture

All fly stocks were maintained on food containing 10% w/v Brewer’s yeast, 8% fructose, 2% polenta and 0.8% agar supplemented with propionic acid and nipagin. Crosses for experiments were performed at 18° (for crosses with temperature inducible gene expression) or 25°. Flies were shifted to 29° after eclosion where relevant.

Male flies were used for all experiments.

The following original fly stocks were used for crosses:

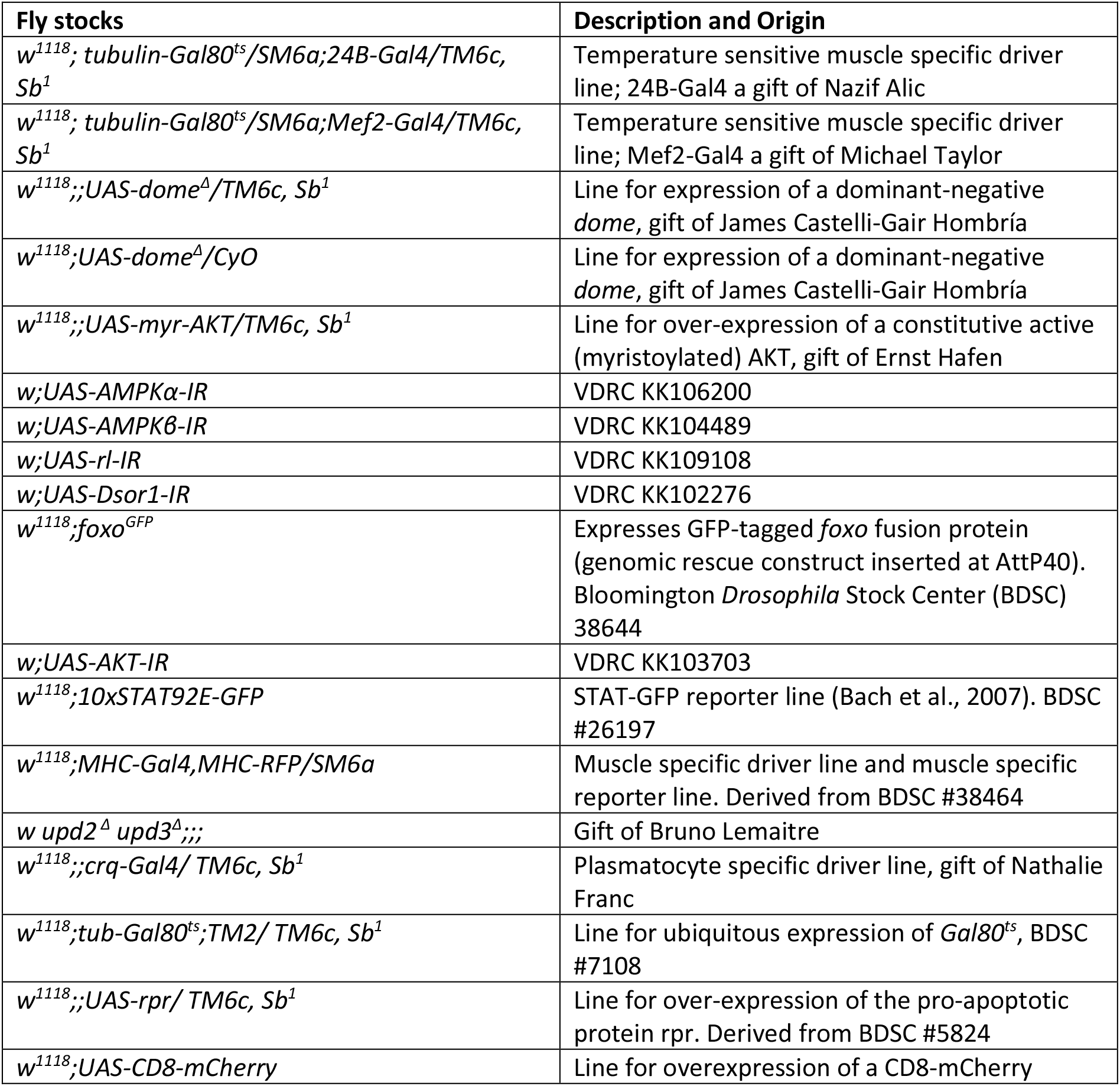

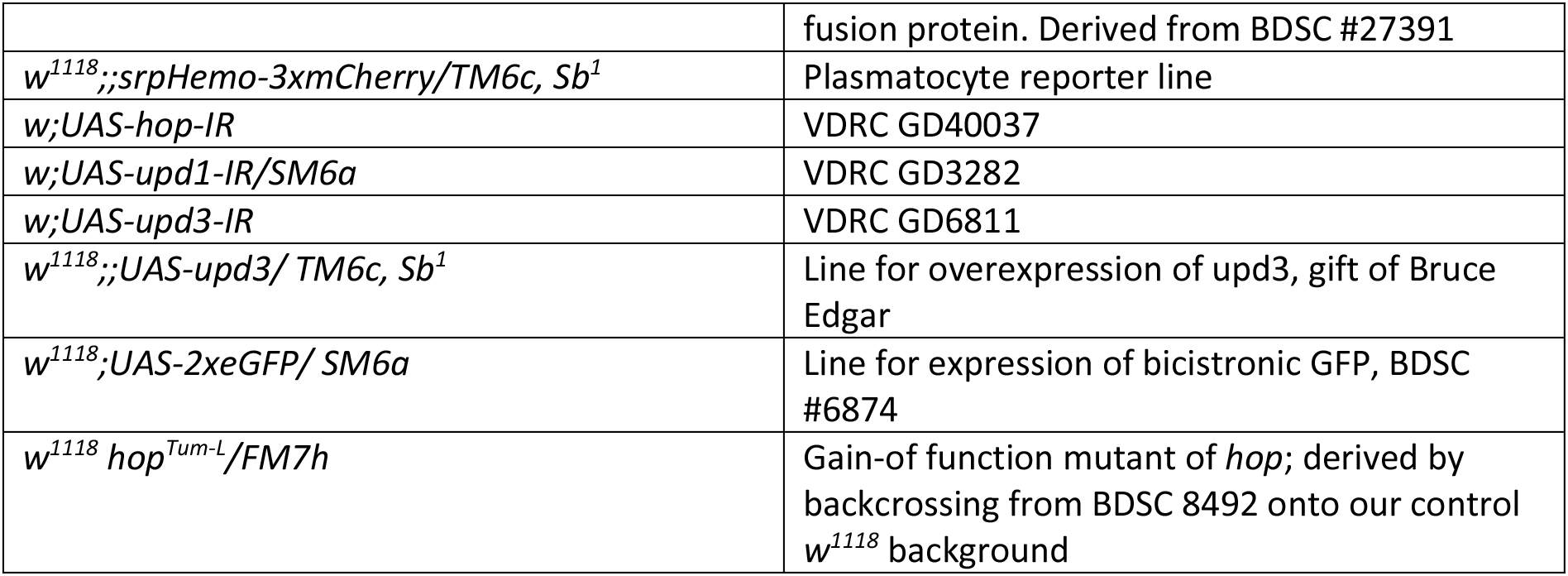

Genotype abbreviations were used for the different experimental flies in this study, in the following table the complete genotypes are indicated:

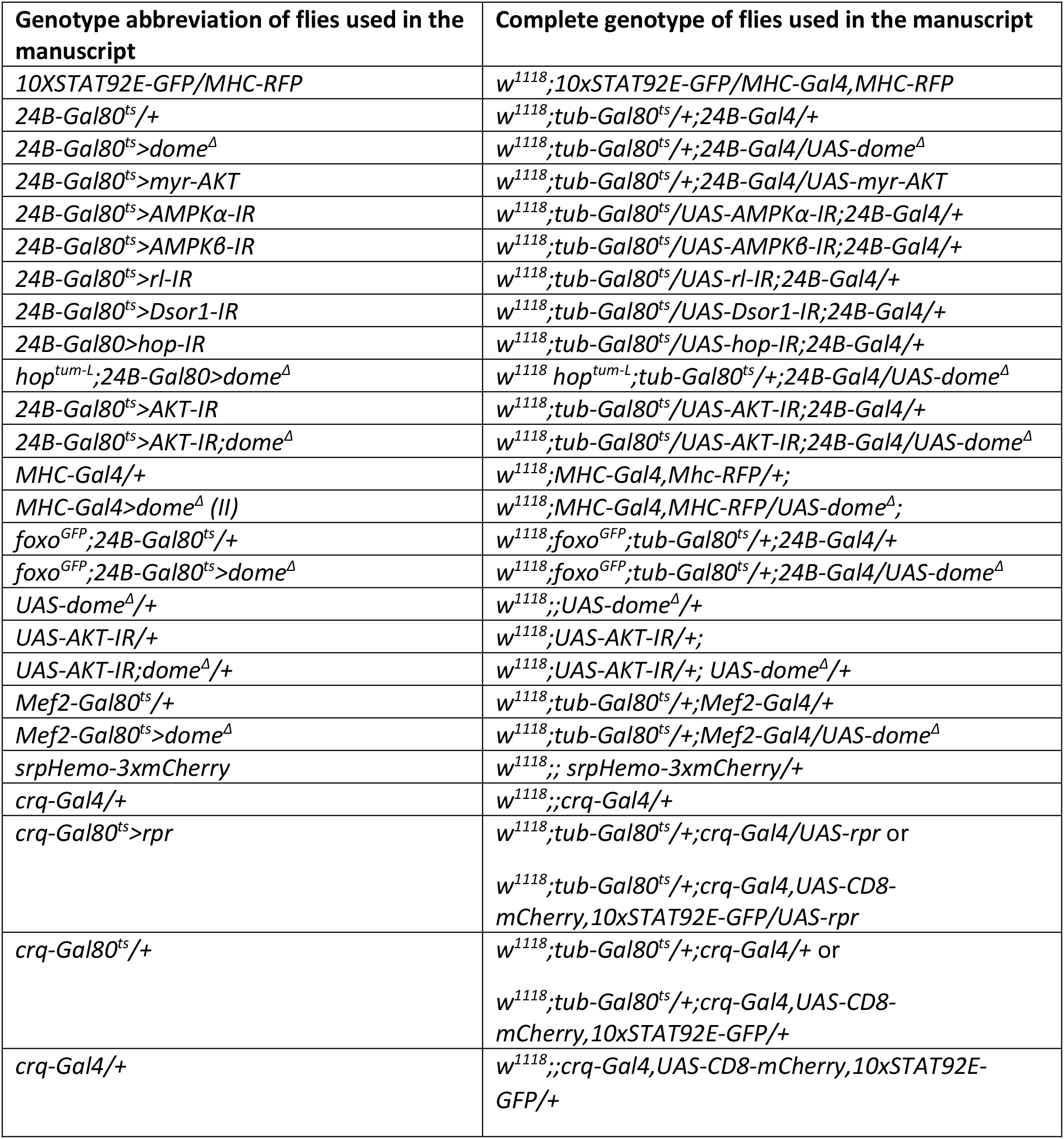

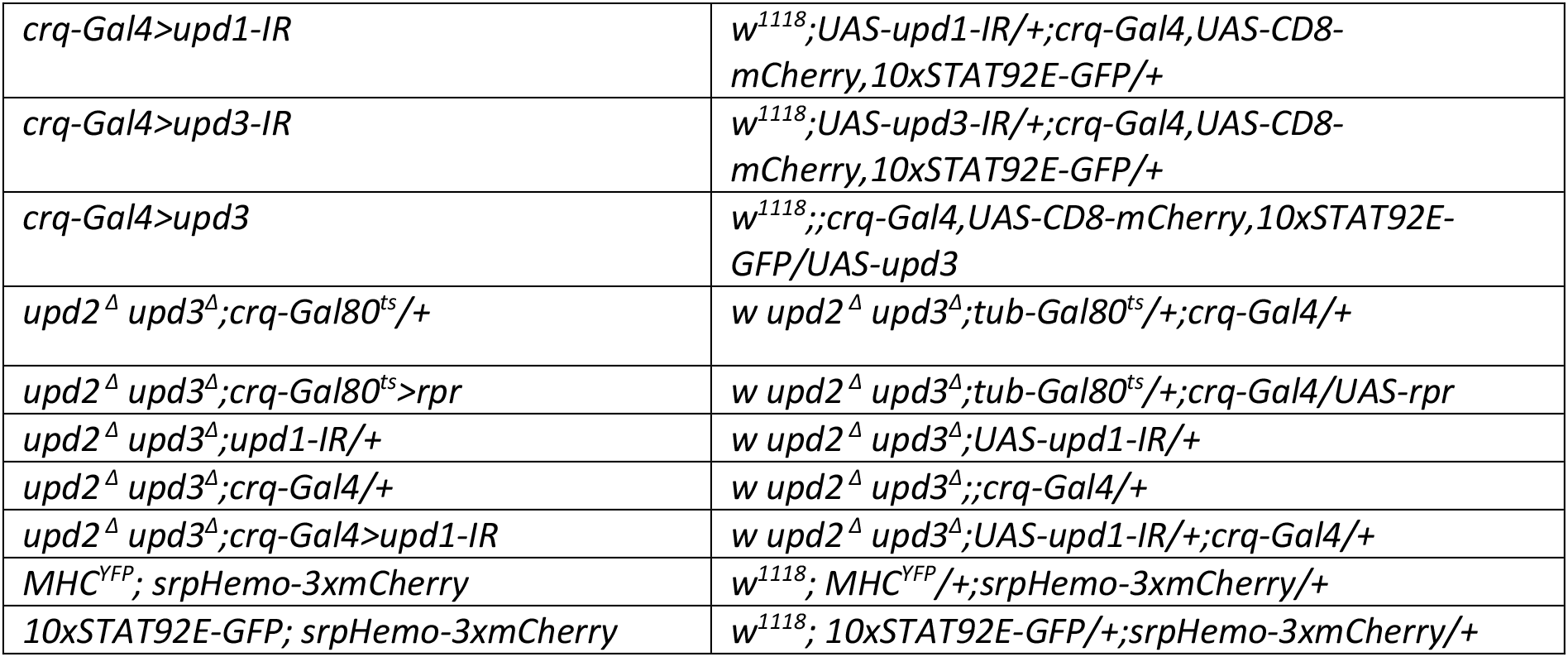

### Lifespan/Survival assays

Male flies were collected after eclosion and groups of 20–40 age-matched flies per genotype were placed together in a vial with fly food (a cohort size of 20 is sufficient to detect a lifespan effect size of about 5% at p=0.05 with 90% confidence). All survival experiments were performed at 29°. Dead flies were counted daily. Vials were kept on their sides to minimize the possibility of death from flies becoming stuck to the food, and flies were moved to fresh food twice per week. Flies were transferred into new vials without CO_2_ anaesthesia.

### Negative Geotaxis Assay/Climbing Assay

Male flies were collected after eclosion and housed for 14 days in age-matched groups of around 20. The assay was performed in the morning, when flies were most active. Flies were transferred without CO_2_ into a fresh empty vial without any food and closed with the open end of another empty vial. Flies were placed under a direct light source and allowed to adapt to the environment for 20 min. Negative geotaxis reflex was induced by tapping the flies to the bottom of the tube and allowing them to climb up for 8 seconds. After 8 seconds the vial was photographed. This test was repeated 3 times per vial with 1 min breaks in between. The height each individual fly had climbed was measured in Image J and the average between all three runs per vial calculated.

### Staining of thorax samples

For immunofluorescent staining of thorax muscles, we anaesthetized flies and removed the head, wings and abdomen from the thorax. Thorax samples were pre-fixed for 1 hour in 4% PFA rotating at room temperature. Thoraces were then halved sagitally with a razor blade and fixed for another 30 minutes rotating at room temperature. Samples were washed with PBS + 0.1% Triton X-100, then blocked for 1 h in 3% bovine serum albumin (BSA) in PBS + 0.1% Triton X-100.

For Lipid-Tox staining, samples were washed with PBS and stained for 2 hours at room temperature with HCS Lipid Tox Deep Red (Thermo Fisher #H34477; 1:200). For Phalloidin labelling, the samples were washed in PBS after fixation and stained for 2 hours at room temperature with Alexa Fluor 488 Phalloidin (Thermo Fisher #A12379, 1:20). Afterwards the samples were washed once with PBS and mounted in Fluoromount-G. All mounted samples were sealed with clear nail polish and stored at 4° until imaging.

### Confocal microscopy

Imaging was performed in the Facility for Imaging by Light Microscopy (FILM) at Imperial College London and in the Institute of Neuropathology in Freiburg. A Leica SP5 and SP8 microscope (Leica) were used for imaging, using either the 10x/NA0.4 objective, or the 20x/NA0.5 objective. Images were acquired with a resolution of either 1024×1024 or 512×512, at a scan speed of 400Hz. Averages from 3-4 line scans were used, sequential scanning was employed where necessary and tile scanning was used in order to image whole flies. For imaging of whole live flies, the flies were anaesthetized with CO_2_ and glued to a coverslip. Flies were kept on ice until imaging. For measuring mean fluorescence intensity, a z-stack of the muscle was performed and the stack was projected in an average intensity projection. Next the area of the muscle tissue analyzed was defined and the mean fluorescent intensity within this area was measured. Images were processed and analysed using Image J.

### RNA isolation and Reverse Transcription

For RNA extraction three whole flies or three thoraces were used per sample. After anaesthetisation, the flies were smashed in 100μl TRIzol (Invitrogen), followed by a chloroform extraction and isopropanol precipitation. The RNA pellet was cleaned with 70% ethanol and finally solubilized in water. After DNase treatment, cDNA synthesis was carried out using the First Strand cDNA Synthesis Kit (Thermo Scientific) and priming with random hexamers (Thermo Scientific). cDNA samples were further diluted and stored at −20° until analysis.

### Quantitative Real-time PCR

Quantitative Real-time PCR was performed with Sensimix SYBR Green no-ROX (Bioline) on a Corbett Rotor-Gene 6000 (Corbett). The cycling conditions used throughout the study were as follows: Hold 95° for 10 min, then 45 cycles of 95° for 15s, 59° for 30s, 72° for 30s, followed by a melting curve. All calculated gene expression values were measured in arbitrary units (au) according to diluted cDNA standards run in each run and for each gene measured. All gene expression values are further normalized to the value of the loading control gene, Rpl1, prior to further analysis.

The following primer sequences have been used in this study:

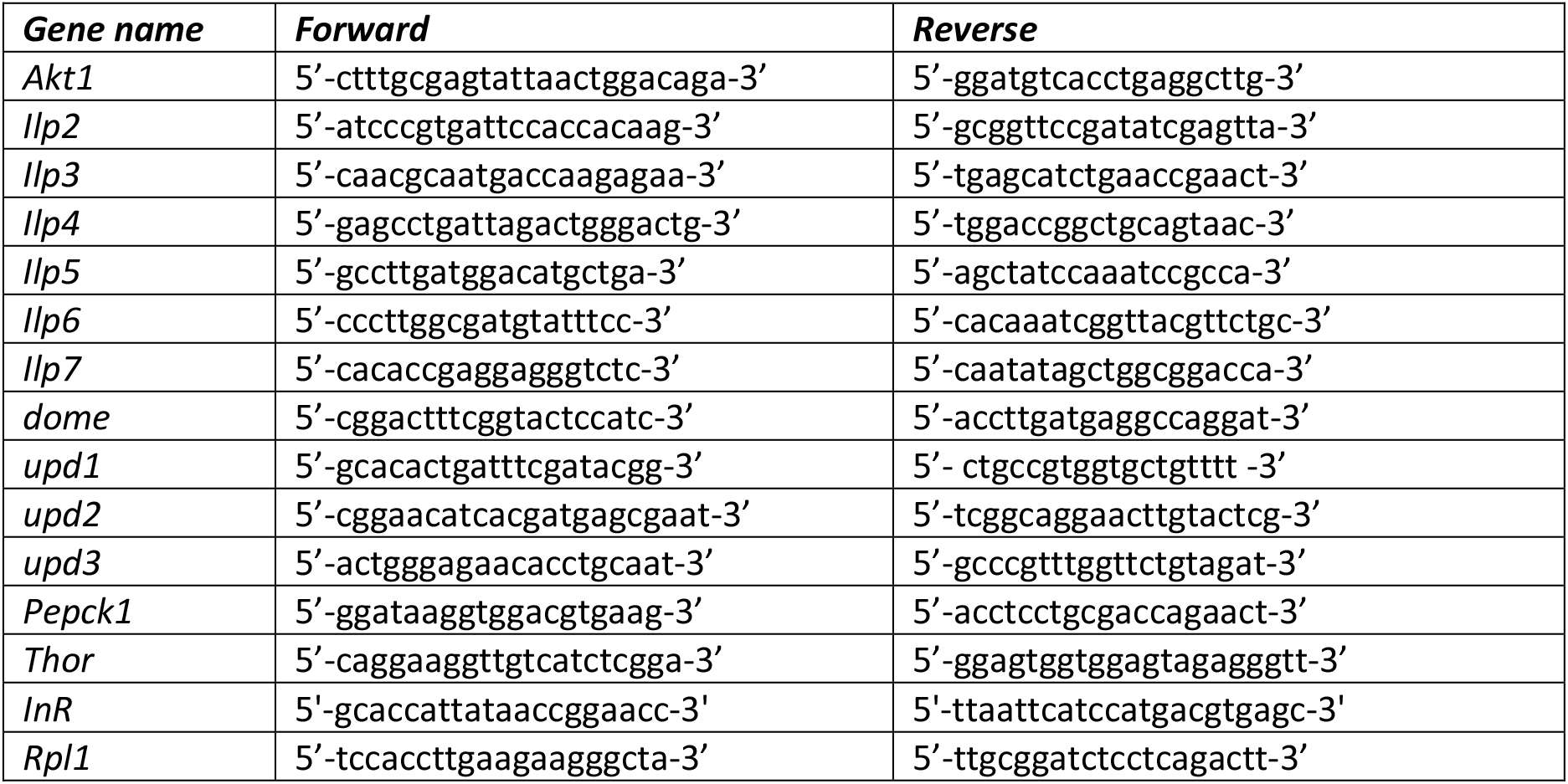

### Smurf Assay

Smurf assays with blue-coloured fly food were performed to analyse gut integrity in different genotypes. Normal fly food, as described above, was supplemented with 0.1% Brilliant Blue FCF (Sigma Aldrich). Experimental flies were placed on the blue-coloured fly food at 9AM and kept on the food for 2 h at 29°. After 2 h the distribution of the dye within the fly was analysed for each individual. Flies without any blue dye were excluded, flies with a blue gut or crop were identified as “non-smurf” and flies which turned completely blue or showed distribution of blue dye outside the gut were classified as “smurf”.

### Western Blot

Dissected legs or thoraces from three flies were used per sample and smashed in 75μl 2x Laemmli loading buffer (100 mM Tris [pH 6.8], 20% glycerol, 4% SDS, 0.2 M DTT). Samples were stored at −80° until analysis. 7.5μl of this lysate were loaded per lane. Blue pre-stained protein standard (11-190kDa) (New England Biolabs) was used. Protein was transferred to nitrocellulose membrane (GE Healthcare). Membrane was blocked in 5% milk in TBST (TBS + 0.1% Tween-20). The following primary antibodies were used: anti-phospho(Ser505)-AKT (Cell Signal Technology (CST) 4054, 1:1,000), anti-AKT (CST 4691, 1:1,000), anti-phospho(Thr172)-AMPKα (CST 2535, 1:1,000), anti-phospho(Thr389)-p70 S6 kinase (CST 9206, 1:1,000), anti-GFP (CST 2956, 1:1,000), anti-phospho-p44/42 MAPK (Erk1/2) (CST 4370, 1:1,000) and anti-α-tubulin (clone 12G10, Developmental Studies Hybridoma Bank, used as an unpurified supernatant at 1:3,000; used as a loading control for all blots). Primary antibodies were diluted in TBST containing 5% BSA and incubated over night at 4°. Secondary antibodies were HRP anti-rabbit IgG (CST 7074, 1:5,000) and HRP anti-mouse IgG (CST 7076, 1:5,000). Proteins were detected with Supersignal West Pico Chemiluminescent Substrate (Thermo Scientific) or Supersignal West Femo Chemiluminescent Substrate (Thermo Scientific) using a LAS-3000 Imager (Fujifilm). Bands were quantified by densitometry using Image J. Quantifications reflect all experiments performed; representative blots from single experiments are shown.

### Thin Layer Chromatography (TLC) for Triglycerides

Groups of 10 flies were used per sample. After CO_2_ anaesthesia the flies were placed in 100μl of ice-cold chloroform:methanol (3:1). Samples were centrifuged for 3 min at 13,000 rpm at 4°, and then flies were smashed with pestles followed by another centrifugation step. A set of standards were prepared using lard (Sainsbury’s) in chloroform:methanol (3:1) for quantification. Samples and standards were loaded onto a silica gel glass plate (Millipore), and a solvent mix of hexane:ethyl ether (4:1) was prepared as mobile phase. Once the solvent front reached the top of the plate, the plate was dried and stained with an oxidising staining reagent containing ceric ammonium heptamolybdate (CAM) (Sigma Aldrich). For visualization of the oxidised bands, plates were baked at 80° for 20 min. Baked plates were imaged with a scanner and triglyceride bands were quantified by densitometry according to the measured standards using Image J.

### Measurement of Glucose, Trehalose and Glycogen

5-7 day old male flies, kept at 29°, were used for the analysis. Flies were starved for 1 hr on 1% agar supplemented with 2% phosphate buffered saline (PBS) at 29° before being manually smashed in 75μl TE + 0.1% Triton X-100 (Sigma Aldrich). 3 flies per sample were used. For measuring thorax, head and abdomen samples, flies were first anaesthetized with CO_2_. Afterwards, they were quickly transferred to 1xPBS and the head was cut off. The guts were carefully removed from thorax and abdomen and thorax were separated from each other. Afterwards the body parts were rinsed with 1xPBS before smashing them in 75μl TE + 0.1% Triton X-100. All samples were incubated at 75° for 20 min and stored at −80°. Samples were thawed prior to measurement and incubated at 65° for 5 min to inactivate fly enzymes. A total of 10μl per sample was loaded for different measurements into flat-bottom 96-well tissue culture plates. Each fly sample was measured four times, first diluted in water for calculation of background fly absorbance, second with glucose reagent (Sentinel Diagnostics) for the measurement of free glucose, third with glucose reagent plus trehalase (Sigma Aldrich) for trehalose measurement, and fourth with glucose reagent plus amyloglucosidase (Sigma Aldrich) for glycogen measurement. Plates were then incubated at 37° for 1 h before reading with a microplate reader (biochrom) at 492 nm. Quantities of glucose, trehalose and glycogen were calculated according to measured standards.

### Respirometry

Respiration in flies was measured using a stop-flow gas-exchange system (Q-Box RP1LP Low Range Respirometer, Qubit Systems, Ontario, Canada, K7M 3L5). Ten flies from each genotype were put into an airtight glass tube and supplied with our standard fly food via a modified pipette tip. Each tube was provided with CO_2_-free air while the ‘spent’ air was concurrently flushed through the system and analysed for its CO_2_ and O_2_ content. All vials with flies were normalized to a control vial with food but no flies inside. In this way, evolved CO_2_ per chamber and consumed O_2_ per chamber were measured for each tube every ~ 44 min (the time required to go through each of the vials in sequence)

### Statistical analysis and handling of data

For real-time quantitative PCR, TLCs, MFI quantification, western blot quantifications and colorimetric measurements for glucose, trehalose and glycogen levels an unpaired t-test was used to calculate statistical significance for all experiments. Respirometer data was analysed with a Mann-Whitney test. Lifespan/ Survival assays, where analysed with the Log-Rank and Wilcoxon test. Stars indicate statistical significance as followed: * p<0.05, ** p<0.01 and *** p<0.001. All statistical tests were performed with Excel or GraphPad Prism software.

All replicates are biological. No outliers were omitted, and all replicates are included in quantitations (including in cases where a single representative experiment is shown). Flies were allocated into experimental groups according to their genotypes. Masking was not used. For survival experiments, typically, the 50% of flies that eclosed first from a given cross were used for an experiment. For smaller-scale experiments, flies were selected randomly from those of a given age and genotype.

## Acknowledgements

We thank the Vienna Drosophila RNAi Center, the Bloomington Drosophila Stock Center, James Castelli-Gair Hombría, Ernst Hafen, Michael Taylor, Dan Hultmark, Nazif Alic, Bruce Edgar, and the FlyTrap collection at Yale University for flies. We are grateful to Rebecca Berdeaux, Günter Fritz, Marco Prinz, Katie Woodcock, Frederic Geissmann, and members of the South Kensington Fly Room for support, discussion and comments. We thank Maria Oberle for technical assistance. Work in the Dionne lab was supported by funding from BBSRC (BB/P000592/1, BB/L020122/2), MRC (MR/L018802/2), and the Wellcome Trust (207467/Z/17/Z). KK was supported by a DFG fellowship. JS was supported by BBSRC/GSK CASE studentship BB/L502169/1. FH was supported by the Neuromac Graduate School of the SFB/TRR167 and the DFG under Germany’s Excellence Strategy (CIBSS-EXC-2189-Project ID 390939984). OG was supported by ERC Starting Grant 337689. The Facility for Imaging by Light Microscopy (FILM) at Imperial College London is part-supported by funding from the Wellcome Trust (104931/Z/14/Z) and BBSRC (BB/L015129/1).

## Author Contributions

MSD and KK designed the project and wrote the manuscript. KK, FH, JS, CMV, PU, AG, DES and JD constructed reagents, performed the experiments and analysed the experimental data.

## Supplemental Information

Figures S1-S4.

**Figure S1.**
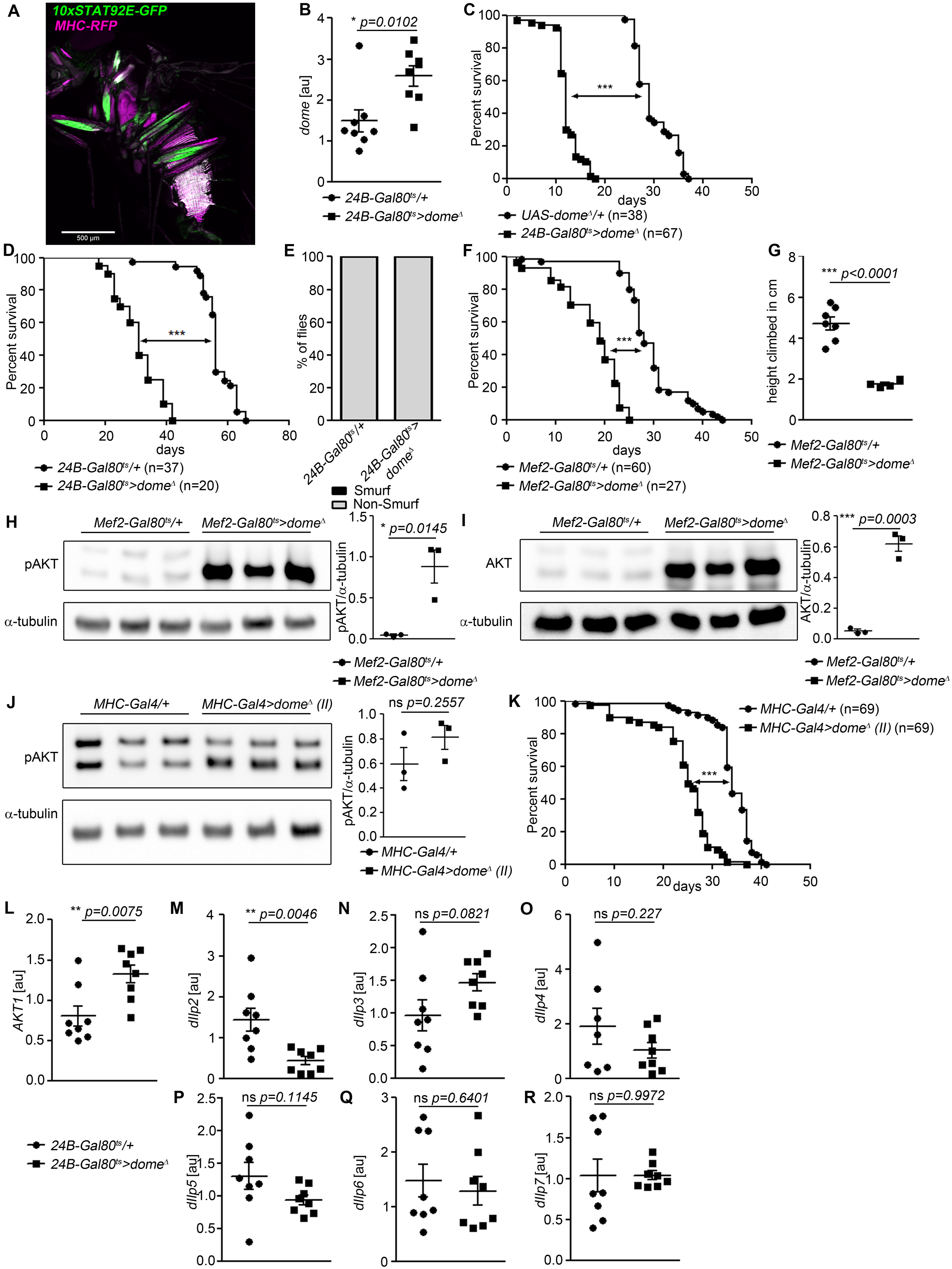
Further characterisation of the requirement for *dome* in adult muscle. (A) STAT activity (*10xSTAT92E-GFP*) and muscle (*MHC-RFP*) colocalize in adult flies. One fly of 6 shown. Scale bar=500μm. (B) *dome* expression by qRT-PCR in thorax samples from 14-day-old *24B-Gal80*^*ts*^*/+* and *24B-Gal80*^*ts*^>*dome*^*Δ*^ flies. (C) Lifespan of *UAS-dome^Δ^/+* and *24B-Gal80*^*ts*^>*dome*^*Δ*^ at 29°; pooled data from two independent experiments shown. Log-Rank test: χ2 =100.8, *** p<0.0001; Wilcoxon test: χ2 =76.2, *** p<0.0001. (D) Lifespan of *24B-Gal80*^*ts*^*/+* and *24B-Gal80*^*ts*^>*dome*^*Δ*^ at 25°; pooled data from two independent experiments shown. Log-Rank test: χ2 =61.83, *** p<0.0001; Wilcoxon test: χ2 =55.18, *** p<0.0001. (E) Smurf assay of 14-day-old *24B-Gal80*^*ts*^*/+* (n=49) and *24B-Gal80*^ts^-*dome*^*Δ*^ flies (n=18). Data pooled from two independent experiments. (F) Lifespan of *Mef2-Gal80^ts^/+* and *Mef2-Gal80*^*ts*^>*dome*^*Δ*^ flies at 29°, pooled from three independent experiments. Log-Rank test: χ2 =86.96, *** p<0.0001; Wilcoxon test: χ2 =78.61, *** p<0.0001. (G) Negative geotaxis assay of 14-day-old *Mef2-Gal80^ts^/+* and *Mef2-Gal80*^*ts*^>*dome*^*Δ*^ flies. Points represent mean climbing height of individual vials analysed (~20 flies/vial), pooled from three independent experiments. (H, I) Western blots of protein from legs of 14-day-old *Mef2-Gal80^ts^/+* and *Mef2-Gal80*^*ts*^>*dome*^*Δ*^ flies. One of three independent experiments is shown. (H) Phospho-AKT. (I) Total AKT. (J) Western blots of Phospho-AKT in leg samples from 14-day-old *MHC-Gal4/+* and *MHC-Gal4>dome^Δ^* (II) flies. One of two independent experiments is shown. (K) Lifespan of *MHC-Gal4/+* and *MHC-Gal4*>*dome*^*Δ*^ (II) flies at 29°, pooled from two independent experiments. Log-Rank test: χ2 =82.9, *** p<0.0001; Wilcoxon test: χ2=58.91, *** p<0.0001. (L-R) Expression by qRT-PCR of *Akt1* and insulin-like peptides in whole fly samples from 14-day-old *24B-Gal80*^*ts*^*/+* and *24B-Gal80*^*ts*^−*dome*^*Δ*^ flies. All transcript levels are normalized to *Rpl1* and shown in arbitrary units [au]. P values in B, G, H-J, L-R from unpaired T-test.

**Figure S2.**
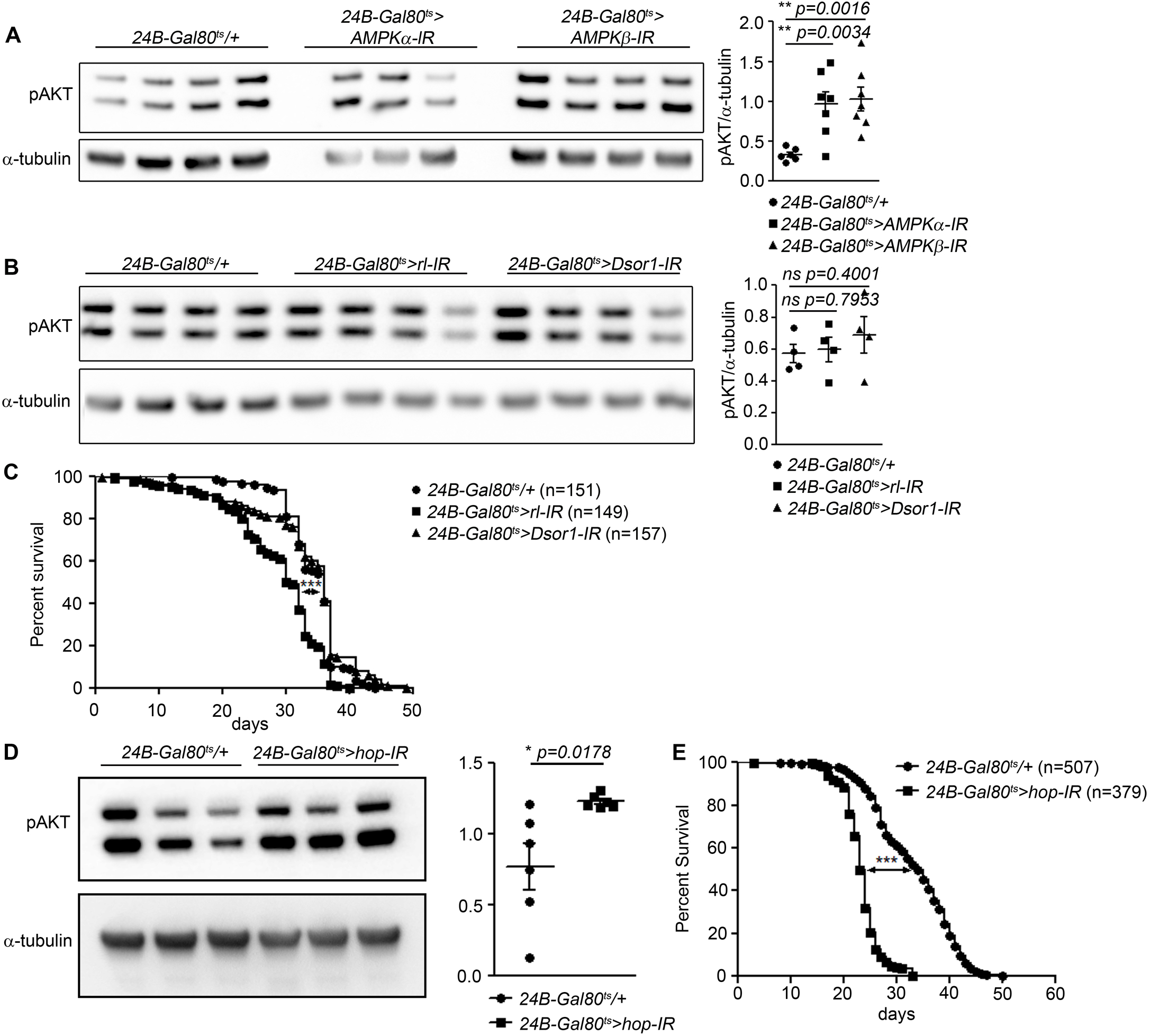
Interactions of *dome* with AMPK, MAPK, and FOXO signaling in adult muscle. (A) Phospho-AKT in leg samples from 14-day-old *24B-Gal80*^*ts*^*/+*, *24B-Gal80^ts^>AMPKα-IR,* and *24B-Gal80^ts^>AMPKβ-IR* flies. One of three independent experiments is shown. (B) Phospho-AKT in leg samples from 14-day-old *24B-Gal80*^*ts*^*/+*, *24B-Gal80^ts^>rl-IR*, and *24B-Gal80^ts^>Dsor1-IR* flies. One of three independent experiments is shown. (C) Lifespan of *24B-Gal80*^*ts*^*/+*, *24B-Gal80^ts^>rl-IR*, and *24B-Gal80^ts^>Dsor1-IR* flies at 29°, pooled from four independent experiments. Log-Rank test (*24B-Gal80*^*ts*^*/+* vs. *24B-Gal80^ts^>rl-IR*): χ2 =60.29, *** p<0.0001; Wilcoxon test (*24B-Gal80*^*ts*^*/+* vs. *24B-Gal80^ts^>rl-IR*): χ2 =58.32, *** p<0.0001; Log-Rank test (*24B-Gal80*^*ts*^*/+* vs. *24B-Gal80^ts^>Dsor1-IR*): χ2 =1.186, ns p=0.2760; Wilcoxon test (*24B-Gal80*^*ts*^*/+* vs. *24B-Gal80^ts^>Dsor1-IR*): χ2 =0.0033, ns p=0.9538. (D) Phospho-AKT in leg samples from 14-day-old *24B-Gal80*^*ts*^*/+* and *24B-Gal80^ts^>hop-IR* flies. One of two independent experiments is shown. (E) Lifespan of *24B-Gal80*^*ts*^*/+* and *24B-Gal80^ts^>hop-IR* flies at 29°, pooled from four independent experiments. Log-Rank test (*24B-Gal80*^*ts*^*/+* vs. *24B-Gal80^ts^>hop-IR*): χ2 =546.4, *** p<0.0001; Wilcoxon test (*24B-Gal80*^*ts*^*/+* vs. *24B-Gal80^ts^>hop-IR*): χ2 =458.1, *** p<0.0001. P values in A, C, E from unpaired T-test.

**Figure S3.**
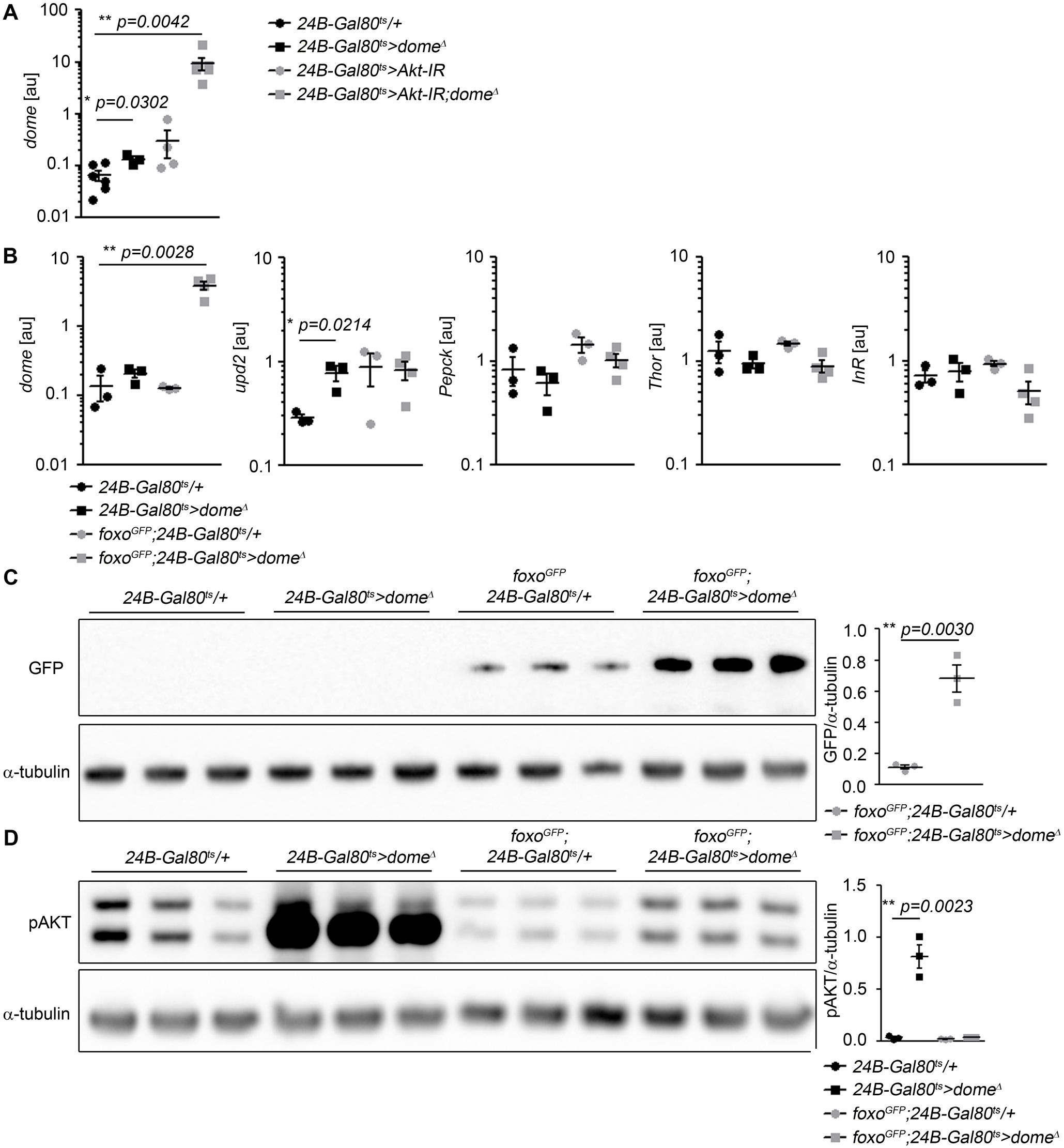
Mutual regulation by AKT, Foxo, and Dome. (A) *dome* expression by qRT-PCR in whole fly samples from 14-day-old *24B-Gal80*^*ts*^*/+*, *24B-Gal80*^*ts*^>*dome*^*Δ*^, *24B-Gal80^ts^>Akt-IR*, and *24B-Gal80*^*ts*^>*Akt-IR;dome*^*Δ*^ flies. (B) Expression by qRT-PCR of *dome, upd2, Pepck, Thor* and *InR* in whole fly samples from 14-day-old *24B-Gal80*^*ts*^*/+*, *24B-Gal80*^*ts*^>*dome*^*Δ*^, *foxo-GFP;24B-Gal80^ts^/+*, and *foxo-GFP;24B-Gal80*^*ts*^>*dome*^*Δ*^ flies. (C) Western blot for GFP to detect the Foxo-GFP fusion protein in leg samples from 14-day-old *24B-Gal80*^*ts*^*/+*, *24B-Gal80*^*ts*^>*dome*^*Δ*^, *foxo-GFP;24B-Gal80^ts^/+*, and *foxo-GFP;24B-Gal80*^*ts*^>*dome*^*Δ*^ flies. (D) Western blot for Phospho-AKT in leg samples from 14-day-old *24B-Gal80*^*ts*^*/+*, *24B-Gal80*^*ts*^>*dome*^*Δ*^, *foxo-GFP;24B-Gal80^ts^/+*, and *foxo-GFP;24B-Gal80*^*ts*^>*dome*^*Δ*^ flies. One of two independent experiments is shown. P values in A-D from unpaired T-test.

**Figure S4.**
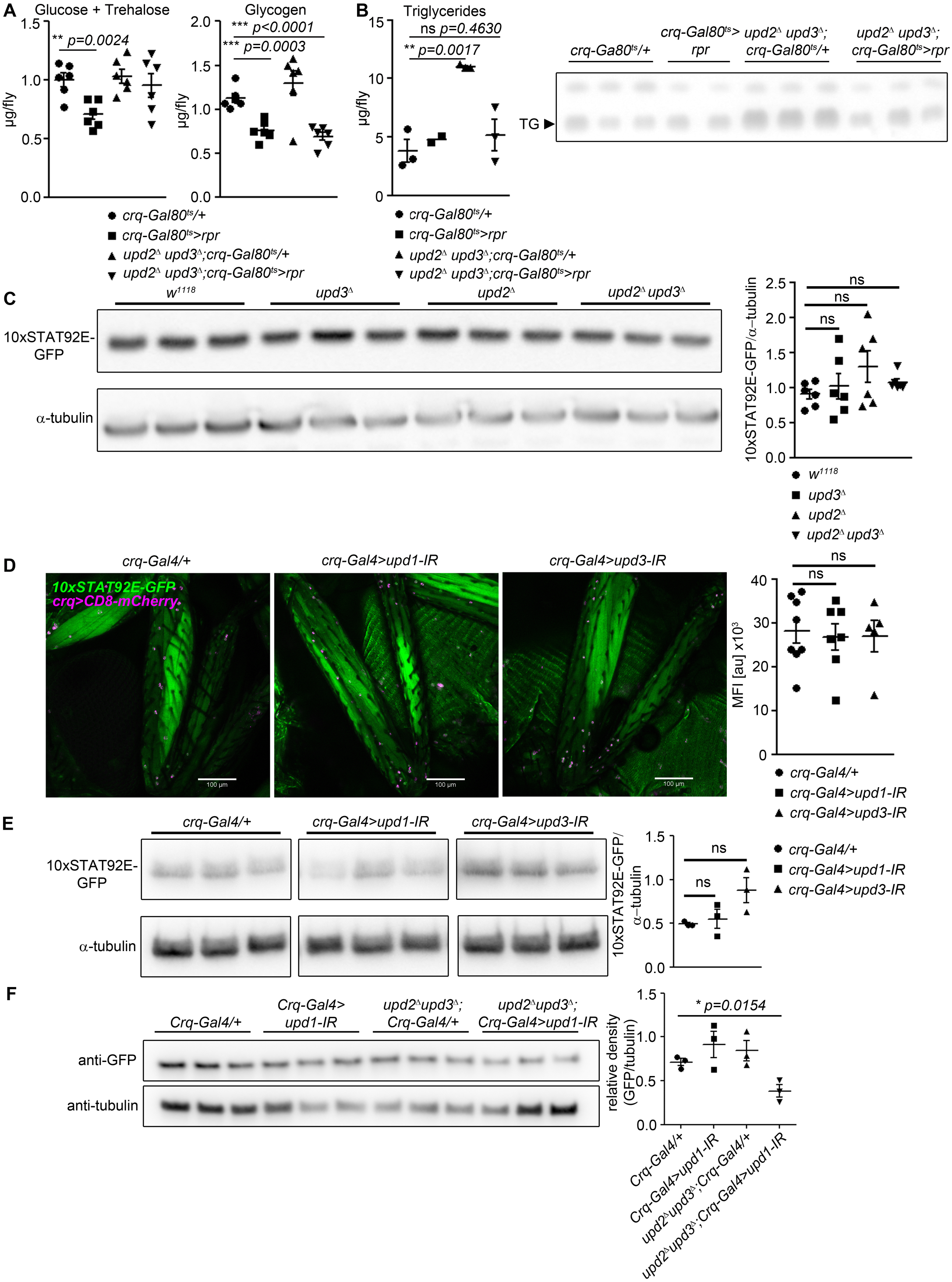
Further characterisation of requirements for specific Upds. (A) Glucose + trehalose and glycogen in 7-day-old *crq-Gal80*^*ts*^*/+*, *crq-Gal80^ts^>rpr*, *upd2^Δ^ upd3^Δ^;crq-Gal80^ts^/+*, and *upd2^Δ^ upd3^Δ^*; *crq-Gal80^ts^>rpr* flies. (B) TLC of triglyceride in 7-day-old *crq-Gal80*^*ts*^*/+*, *crq-Gal80^ts^>rpr*, *upd2^Δ^ upd3^Δ^;crq-Gal80^ts^/+*, and *upd2^Δ^ upd3^Δ^;crq-Gal80^ts^>rpr* flies, n=2-3 samples per genotype. (C) Western blot analysis of STAT-driven GFP in legs from 7-day-old *w^1118^*, *upd3^Δ^*, *upd2^Δ^*, and *upd2*^*Δ*^ *upd3*^*Δ*^ flies. One representative experiment of two is shown. (D) STAT activity and plasmatocytes in legs from 7-day-old control (*crq-Gal4/+*), *upd1* knockdown (*crq-Gal4>upd1-IR*), and *upd3* knockdown (*crq-Gal4>upd3-IR*) flies. One representative fly is shown of 5-7 imaged for each genotype. Scale bar=100μm. Mean fluorescence intensity (MFI) is shown for all flies imaged. (E) Western blot analysis of STAT-driven GFP in legs from 7-day-old control (*crq-Gal4/+*), *upd1* knockdown (*crq-Gal4>upd1-IR*), and *upd3* knockdown (*crq-Gal4>upd3-IR*) flies. One of two independent experiments is shown. (F) Western blot analysis of STAT-driven GFP in thorax from 7-day-old *crq-Gal4/+*, crq-Gal4>*upd1*, *upd2^Δ^upd3^Δ^;crq-Gal4/+* and *upd2^Δ^upd3^Δ^;crq-Gal4>upd1-IR* flies. P values in A-F from unpaired T-test.

